# Scaling up submerged fermentation process of *Aspergillus fumigatus* mga for an efficient metabolite production (xylanase)

**DOI:** 10.1101/2022.07.17.500358

**Authors:** Maroua Gares, Serge Hiligsmann, Noreddine Kacem Chaouche

## Abstract

Fungal metabolites production at an industrial scale requires a sufficient yield at relatively low cost. Indeed, the scaling-up process is one of the main bottlenecks in the fermentation procedure; the reproduction of the best conditions achieved in small bio-reactors when transferring them to a much larger fermentation scale is near impossible.

The present study seeks to study the fermentation profile of *Aspergillus fumigatus* fungi, in order to spot it logarithmic phase using xylose as substrate in different volumes; 250 mL flasks, 2 L and 20 L bioreactors, before conducting further experiments for an efficient secondary metabolite production (xylanase). The agitation speed impact on the morphological changing of this fungi was also studied.

## 1. Introduction

Filamentous fungi have been cultivated for thousands of years. Currently, antibiotics, enzymes, organic acid and additional pharmaceutical products are carried out using metabolites derived from these microorganisms [1]. The use of modern tools of biotechnology and a convenient fermentation physiology are needed to achieve successful large-scale production [2]. However, due to the growing interest in this field, scientists have sought biotechnological alternatives such as submerged fermentation (SmF) with free mycelium, solid-state fermentation (SSF) with the development of mycelium in the absence of free water on a solid substrate, generally agricultural by-products, and also biofilm formation with the development of mycelium on an inert support. Among these three strategies, SmF is the most widely used in the industry; metabolite production is mostly carried out under submerged conditions due to a simpler down-streaming process and the best control over the various parameters [3]. However, one of the major expenses in current enzyme production on a large scale is the sugar cost; microbial xylanases are induced mainly by xylans, which makes the cost of enzyme production very high [4].

Filamentous fungi are well-known producers of xylanase during fungi’ secondary metabolite process. On an industrial scale, xylanases are produced mainly by *Aspergillus* and *Trichoderma spp*. in SSF [5]. *Aspergillus fumigatus* is known as a suitable fungus for the production of heterologous proteins. High enzyme production rates justify the increasing interest in this micro-organism despite its pathogenicity. Indeed, extremely efficient CRISPR-mediated genome editing was shown [6]. Therefore, deletion of genes encoding enzymes of melanin biosynthesis, encoding transcription factors triggering production of secondary metabolites, i.e. gliotoxin, or encoding extracellular proteases and siderophore synthesizing enzymes might result in non-pathogenic strains [7].

This paper describes the fermentation behavior of *A. fumigatus* in submerged cultures scale-up, in order to define its logarithmic phase for further exploitation e.g. large production of metabolites of interest as xylanase.

## 2. Materials and methods

### Scale-up processing using xylose substrate; batch submerged fermentation in 250 mL shake flasks, 2 and 20 L bioreactor

Submerged fermentations were conducted to explore the consumption of xylose using *A. fumigatus* strain. These experiments were carried out in order to test the ability of this fungus to develop in a submerged medium based on simple sugars in different volumes and also to study its fermentation profile.

#### Biomass Growth and pH

The culture broth was separated from the mycelium by filtration using a vacuum pump and filter paper (Whatman N° .1); the mycelium was dried in an oven at 60 °C until constant weight according to AOAC standards. Mycelial biomass concentration was determined by grams of dry weight per liter of liquid culture (g/L).

During the fermentation period, samples were collected, and subsequently the pH was measured with a pH meter for the flask cultures and a pH sensor for the bioreactor fermentations.

#### Xylose quantification

Samples of 5 mL collected each day of fermentation were filtrated and centrifuged at 14,000× g for 15 min. The supernatant obtained was separated from the residual biomass at 4 °C and finally, the soluble molecule concentration (xylose) was determined by HPLC-RID and expressed in g/L.

#### Preparation of standard inoculum for <u>A. fumigatus</u> fermentation in shake flask culture

Spore suspension was prepared from five-day-old cultures of *A. fumigatus* on potato dextrose agar (PDA) at 30°C. Spores were harvested by adding 10 mL of sterile distilled water containing 1% (v/v) tween 80, then collected in sterile flasks to be used as inoculum for further cultures. The spore concentration was adjusted to 1× 10^6^ spores/mL using a counting chamber (Thomas REF 06 106 10 MARIENFELD, Germany) [8]. Then, the culture flasks with a working volume of 100 mL medium were prepared as the following (gram per liter): 10 xylose, 10 peptone, 10 yeast extract in a 250 mL culture flask. The initial medium culture pH was 6.5 without adjustment. Then, the culture medium was sterilized at 121 °C for 20 minutes before inoculation.

One mL volume of fresh conidia suspension was inoculated into the 250-mL Erlenmeyer flasks and finally incubated in a shaking incubator (200 rpm) (Ecotron, INFORS HT, Switzerland) at 30°C.

#### Preparation of standard inoculum for <u>A. fumigatus</u> fermentation in 2 L and 20 L bioreactors

To perform the fermentation scale up in 2 L and 20 L bioreactors, a standard inoculum size of 1×10^6^ spores/mL was inoculated into appropriate corning Erlenmeyer flasks with 150 mL and 1.5L working volume respectively and incubated for 12 h at 120 rpm and 30°C. The flasks transfer caps have two ports. One port ends in a 0.2 μm filter, and the other port is connected with a dip tube that reaches all the way to the bottom of the flask for an easy aseptic transfer of the pre-culture.

The culture medium for both bioreactors was prepared as described in the previous part, the carbon source was prepared and autoclaved separately from other medium compositions. A temperature of 30 °C was adjusted and the inoculation process started when the adjusted temperature was reached. The incubation period was stopped when the substrate was completely consumed (stationary – decline phase).

Before inoculation, the corresponding vessels and important accessory elements of the fermenter were sterilized. The 02L bioreactor sterilization was conducted in a proper laboratory autoclave, the 20 L is a sterilize-in-place bioreactor. The pO2 and pH sensors were calibrated before inoculation; the pre-cultures were transferred with peristaltic pump into the 02 L (DCI-Biolafitte, US) and 20 L (Sartorius Bbraun, US) bioreactors respectively. The fermenters were agitated and aerated at appropriate rates. Antifoam solution was used when necessary. Samples from pre-culture and main-culture were tested for contamination.

#### *Agitation speed impact on the fermentation process and morphological changing of <u>A. fumigatus</u>*; small scale study

Cultivation was carried out with a 2 L bioreactor with the same submerged medium and pre-culture concentration at an agitation speed of 200 rpm. After 24 hours, the stirring speed was increased. Fungal mycelia samples were harvested from various agitation speeds (350, 500 and 600 rpm). 1 mL of these samples were further inoculated into test tubes containing the same liquid medium but with a xylose concentration of (2g/L), and incubated for 5 days. The substrate consumption was analyzed and the morphological change of samples was viewed under a light microscope.

## 3. Results and discussion

### Scale-up processing; batch SmF in 250 mL shake flasks, 2 L and 20 L bioreactor

#### Submerged batch fermentation in shake flasks

Many studies have reported that the best inducers for xylanolytic enzyme production in *Aspergillus* spp., are xylose, xylan and crude xylan-containing substrates [9]. In the current study, xylose was used as the only carbon source to study *A. fumigatus* fermentation profile and to spot its logarithmic phase. The growth profile of dry mycelial cell mass and substrate concentration were presented in (Figure 4 A). The substrate was almost completely consumed after 96 hours of fermentation accompanied with a maximal biomass production (11.81 g/L); the exponential phase was recorded between 24 and 48 hours of fermentation.

The stationary phase was observed between 48 h and 96 h after the biomass decreased rapidly due to the negative effect.

Regarding the pH; it dropped from 6.6 to 5.89 when the biomass reached its maximal production and consumed almost all the substrate. This decrease in the pH may be due to the production of metabolites derived from xylose consumption. These types of fungal strains produce secondary metabolites such as organic acids and enzymes (e.g., xylanase, cellulases, pectinases, proteases, amylases etc.,) [10,11]. The subsequent increase in pH after 96 hours of fermentation, correlates with the depletion of substrate in the medium. Causing an accumulation of nitrogenous wastes, which lead to an increase in the pH (7.04)[12].

#### *Agitation speed impact on the fermentation process and morphological changing of* A. fumigatus

In submerged culture, filamentous fungi show diverse macro-morphology such as hyphal pellets (Figure 2). These mycelial aggregates interfere with the substrate uptake, causing metabolic limitations and decreasing production efficiency. This research study prevented these pellets formation by three steps: (1) the inoculation of the pre-culture with concentrated conidia and decreasing its incubation time, (2) short term culture and (3) high agitation velocity. Achieving a homogeneous medium in SmF can result in efficient mass transfer and a high quality and quantity of production. Many studies have sought optimal conditions for agitation, because mycelial damage at high stirrer speeds or power inputs can limit the capability and volumetric productivity of a fermenter [13]. According to Ghobadi et al., [13], high agitation can cause a huge effect on microorganisms such as cell damage, morphological changes, and variations in growth rate and product formation. For each submerged culture, the optimal conditions for agitation depend partly on the resistance of hyphae to the mechanical forces and also on its physiological state [14].

In the present study, agitation speed impact on *A. fumigatus* morphology was found to accelerate substrate consumption. However, the fermentation process is slightly reduced with 650 rpm samples. Macroscopic morphology at the end of the incubation period showed that hyphae were not attached to the fermenter as in (Figure 1 A), but forming free micro-pellets (Figure 1 B and C). Indeed, in fine-dispersed mycelia, each hyphal filament is reached by the convection of the stirred medium [7].

**Figure 1.**
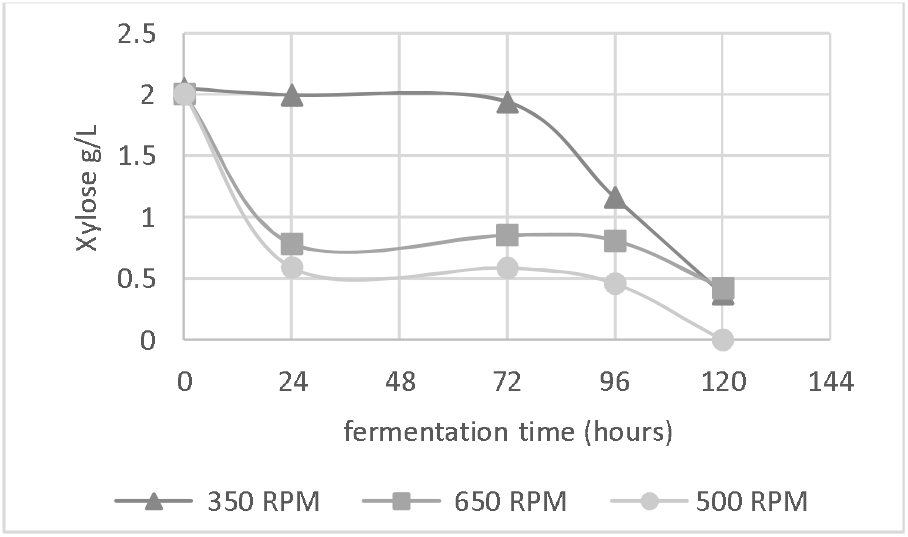
Agitation speed impact on the fermentation profile of *A. fumigatus*

**Figure 2.**
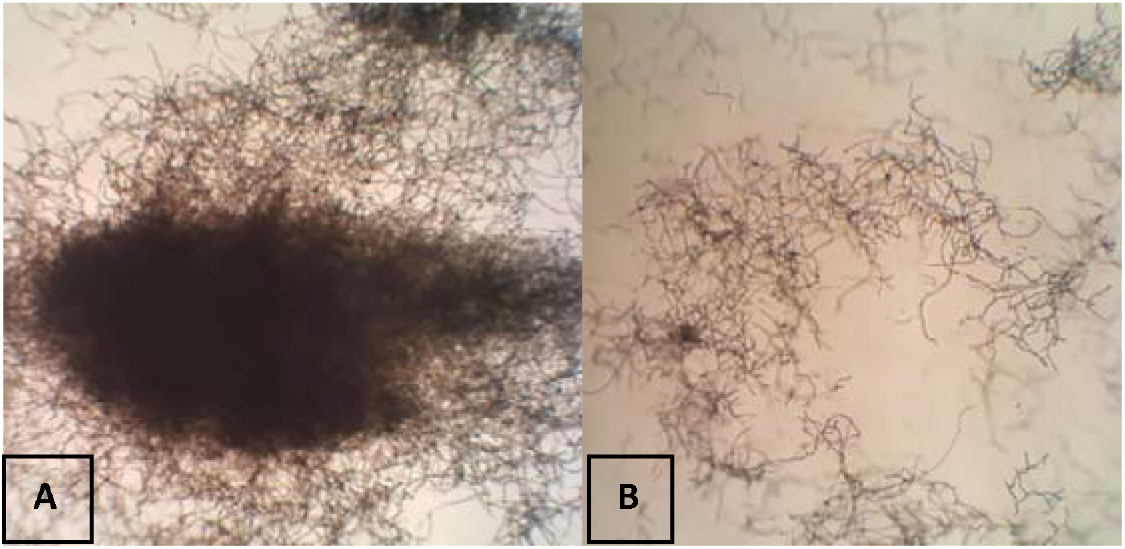
microscopic morphology of *A. fumigatus* flocs (pellets) in stirred vessel cultivation at 200 rpm (A) and “shear effect” at 650 rpm (B)

**Figure 3.**
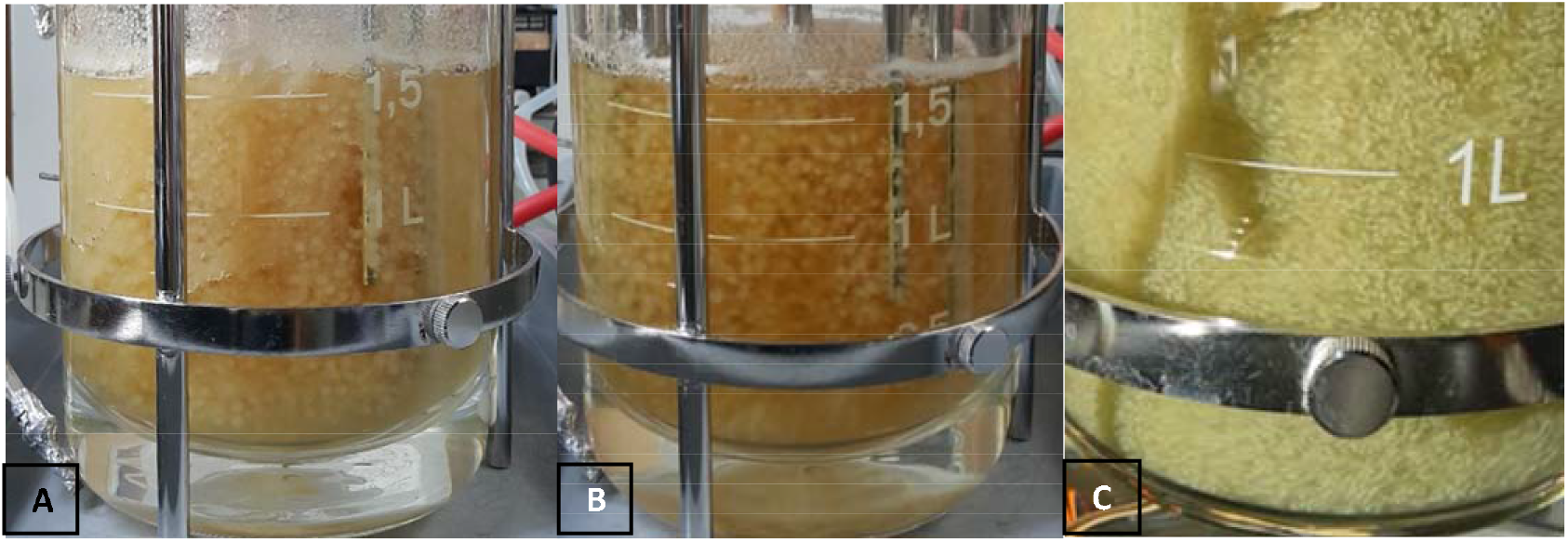
Macroscopic morphology of *A. fumigatus* flocs (pellets) in stirred 2L bioreactor at the end of cultivation process, at 200 rpm (A), 350 rpm (B) and 500 rpm (C)

**Figure 4.**
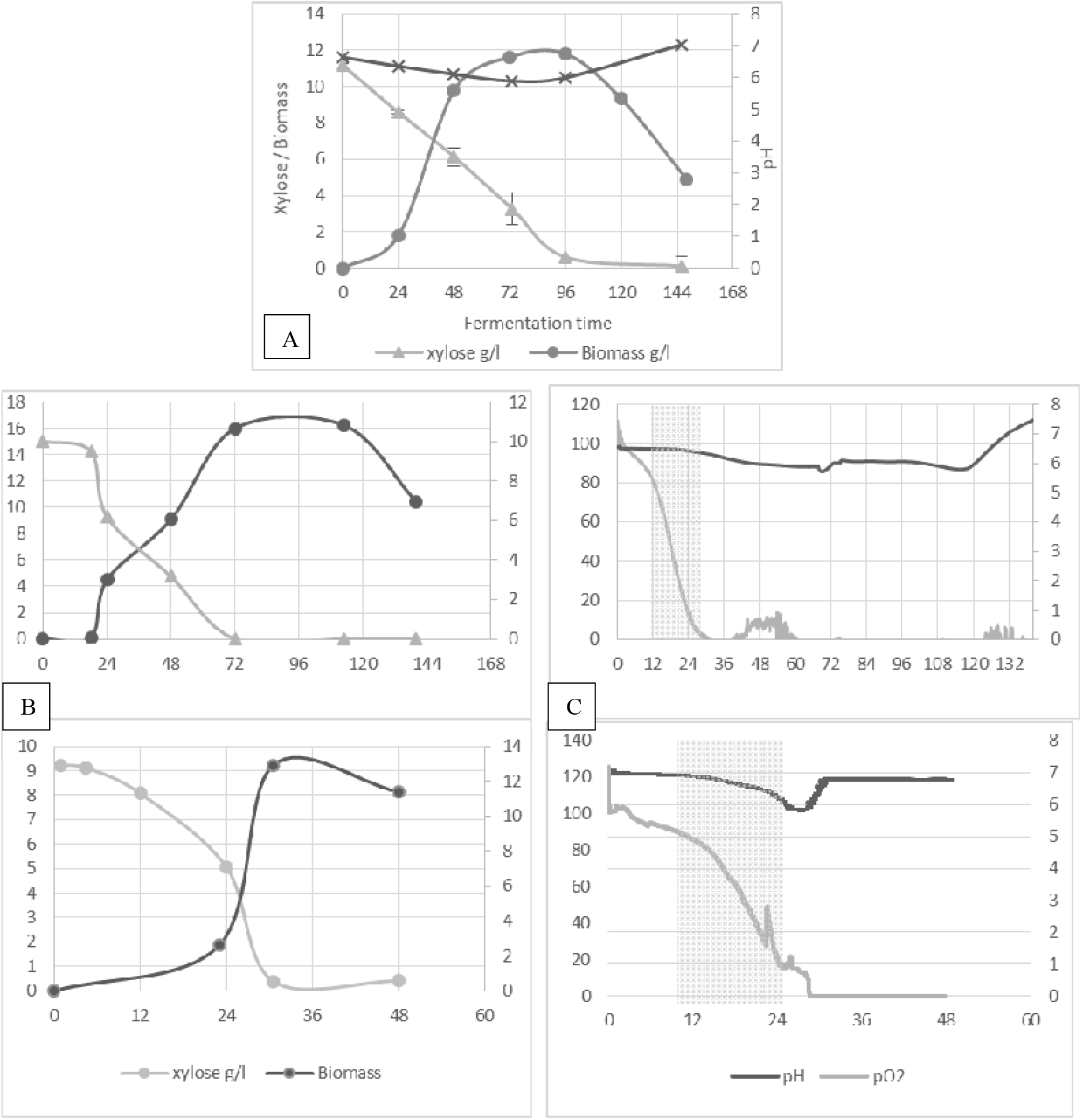
Fermentation profile of *Aspergillus fumigatus* under submerged fermentation in 250 mL flasks (A), 2 L bioreactor (B) and 20 L bioreactor (C).

In macro-pellets, gas exchange as well as transport of substrates and products is hindered by pseudo-tissue. This may be the cause of the exponential phase delay for 350 rpm samples.

#### Submerged fermentation in 02 and 20 L batch bioreactors

In batch cultures, all concentrations change minute by minute. In order to minimize diffusion barriers, high conidia concentrations were used as inoculum for the pre-culture. The short fermentation time avoided macro-pellet formation. Furthermore, in this kind of cultures, oxygen limitation is a very important aspect to be taken into account due to the high viscosity of the fungal broth that results in a much lower gas-liquid mass transfer coefficient (kLa) in comparison to unicellular microorganism fermentation [15].

Indeed, oxygen limitation is a common cause of low level production of aerobic products like enzymes. To circumvent this limitation, Fratebianchi et al., [16] controlled the effects of airflow and agitation speed to enhance polygalacturonase production in SmF by *Aspergillus sojae*. And found that agitation of 600 rpm and cascading airflow from 1 up to 1.7 vvm, fixed the oxygen limitation problem, therefore, maximum enzyme production was achieved. Same study reported that higher agitation speed causes microorganisms stress leading to a decrease in the enzymatic production. A corresponding reaction was previously reported regarding recombinant protein production by *A. niger* in SmF bioreactors [17,18]. Bandaipeth and Prasertsan [19] reported that increasing the airflow is a better choice than increasing agitation speed in order to get high kLa values and avoid microorganisms stress. Another study conducted by Bakri et al., [20] recorded a maximum xylanase activity at agitation speed of 200 rpm and aeration rate of 1.0 vvm. Although, in this agitation speed, macro-pellets formation is inevitable.

In aerobic bioprocesses, oxygen is a key substrate; due to its low solubility in liquid broths, a continuous supply is needed. Therefore, the oxygen transfer rate (OTR) must be defined, and if possible predicted to achieve an optimum design operation and scale-up of bioreactors [21]. Agitation speed and aeration rate in our study were set in a way to maintain kLa constant during scale□up from 2 to 20 L.

Submerged batch cultures are commonly characterized according to the classical growth curve in textbooks (i.e., adaptation, acceleration, exponential, stationary, and declining phase). Although, online respirometry proved to be a powerful tool to distinguish growth phases and revealed more physiological states than expected from the mere biomass curve [22]. In the current research, the monitoring system allows continuous tracking of pH and PO2 in every 05 and 04 min for 02L and 20L bioreactors respectively, which makes the analysis more efficient.

According to the PO_2_ curve (figure 5 B and C), the logarithmic phase of the both 2L and 20L bioreactors was recorded between almost 10 and 24 h. The growth profile of dry mycelial cell mass and substrate concentration were also presented in Figure 5 (C and D). The substrate was completely consumed after 72 hours of fermentation in the 2 L bioreactor and only 30 hours in the 20L bioreactor, the maximal biomass production of 16.22 g/L and 12.9 g/L was reached after 72 and 30 hours of fermentation for the bioreactor of 2L and 20 L respectively.

The pH of the culture media followed the same trajectory as that of the flask culture; it dropped also to almost 5.7 in the both bioreactor cultures when the biomass reached its maximal production, and then, it started to increase at the end of the fermentations.

## Data availability

All data generated or analysed during this study are included in this published article

